# Machine Learning-Enabled Raman Spectroscopy for Process Analytical Technology and Real-Time Release Testing in Bioprocess Manufacturing: A Comparative Predictive Modeling Study

**DOI:** 10.64898/2026.07.24.740653

**Authors:** S. Patel, V. Patel

## Abstract

Analytical technologies that can provide quick, precise, and continuous information regarding process performance are necessary for the development of biopharmaceutical manufacturing. Conventional bioprocess monitoring is largely dependent on laboratory-based data and offline sampling, which can restrict process management and cause delays in decision-making. This study develops a machine learning-enabled Raman spectroscopy framework for Process Analytical Technology (PAT) and Real-Time Release Testing (RTRT) applications in bioprocess manufacturing. Five predictive modeling techniques—Partial Least Squares (PLS) regression, Support Vector Regression (SVR), Random Forest, Extreme Gradient Boosting (XGBoost), and Neural Networks—were used to analyze Raman spectral data from an Escherichia coli fermentation dataset. The models were assessed using the coefficient of determination (R²), root mean square error (RMSE), and mean absolute error (MAE) to predict two crucial fermentation parameters: the concentrations of glucose and acetate. The superior performance of PLS regression for glucose prediction and the improved prediction accuracy of XGBoost for acetate concentration demonstrated the importance of selecting modeling techniques based on biological complexity. Explainable artificial intelligence using SHAP analysis was incorporated to improve model transparency by identifying Raman spectral regions contributing to predictions. The suggested architecture shows how Raman spectroscopy and machine learning can be combined to assist automated process monitoring, enhance process comprehension, and hasten the implementation of real-time quality judgments in next-generation biomanufacturing.

**Graphical Abstract:** 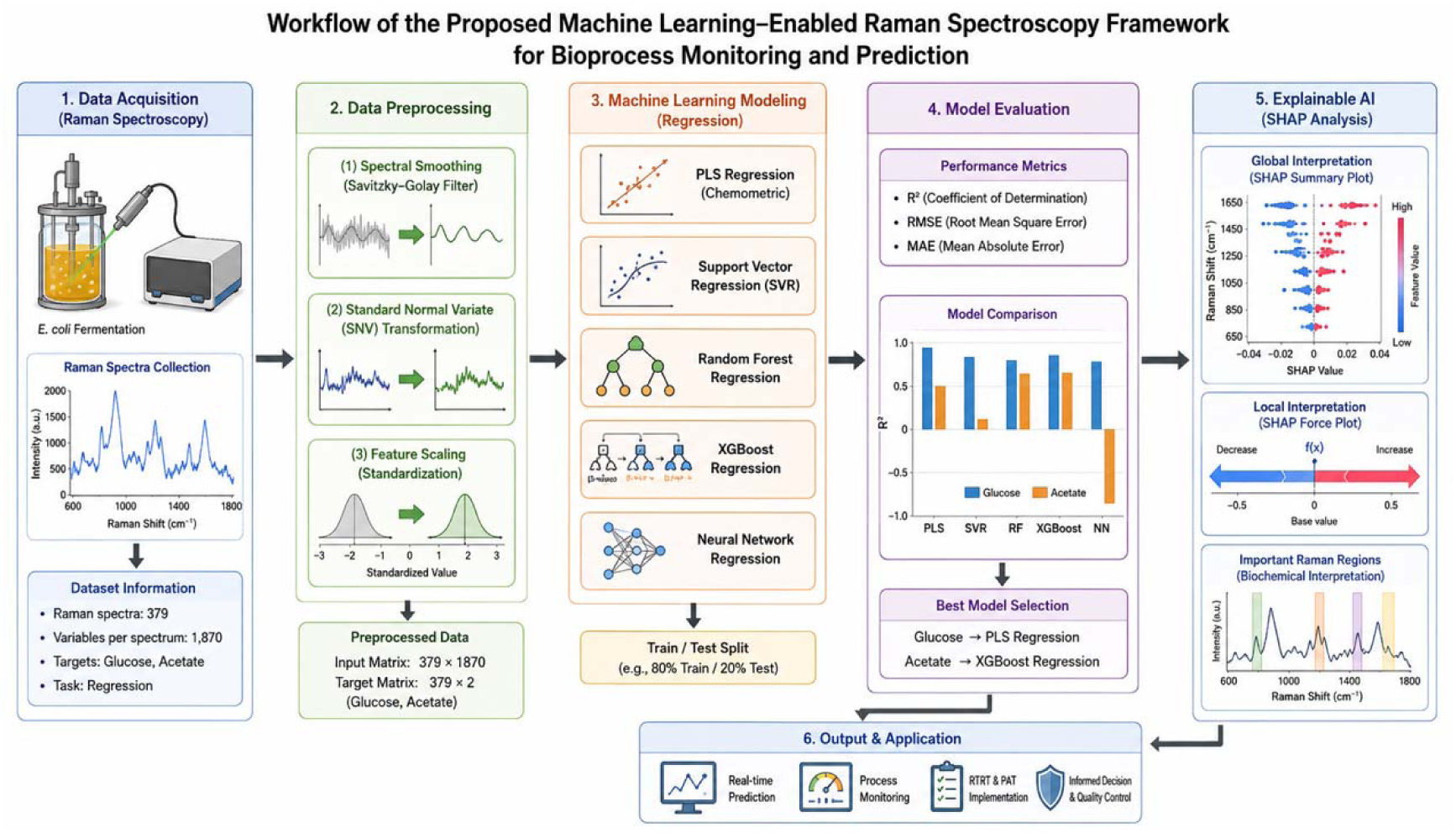

Overall workflow of the Raman spectroscopy-based machine learning framework for PAT and RTRT implementation. Raman spectra collected from *E. coli* fermentation were preprocessed and analyzed using multiple machine learning algorithms for the prediction of glucose and acetate concentrations. Model performance evaluation and SHAP-based explainable AI analysis enabled the identification of important spectral features for real-time bioprocess monitoring.

**Highlights:**

- Developed a Raman spectroscopy-based machine learning framework for real-time monitoring of critical bioprocess parameters.
- Compared traditional chemometric modeling (PLS regression) with advanced machine learning approaches, including SVR, Random Forest, XGBoost, and neural networks.
- Showed that the biochemical target affects the model’s performance, with XGBoost improving acetate prediction and PLS offering better glucose prediction.
- Integrated explainable artificial intelligence to identify Raman spectral regions contributing to bioprocess predictions.
- Established a pathway toward interpretable Raman-based Process Analytical Technology (PAT) and Real-Time Release Testing (RTRT) implementation.

## 1. Introduction

A significant shift toward automated, continuous, and data-driven production systems is taking place in the biopharmaceutical sector. Instead of depending on postponed offline laboratory testing, modern manufacturing methods increasingly call for analytical systems that can monitor crucial process parameters in real time. Through real-time measurement, process comprehension, and enhanced control techniques, Process Analytical Technology (PAT) was developed to enhance pharmaceutical manufacturing [1].

Conventional bioprocess monitoring techniques often rely on sample removal and subsequent laboratory analysis. These techniques are time-consuming and could miss the quick biological changes that take place during fermentation, even if they offer accurate readings. Because it allows for non-destructive, quick, and in-line monitoring of biological systems without needing much sample preparation, Raman spectroscopy has become a popular PAT technique [2], [3].

Raman spectroscopy provides molecular information by measuring vibrational characteristics of chemical bonds within biological samples. In fermentation processes, Raman spectra contain information related to metabolites, nutrients, and cellular conditions. Previous studies have demonstrated the capability of Raman spectroscopy for monitoring glucose, lactate, biomass, and other critical process variables in microbial and mammalian cell cultures [4].

However, the correct interpretation of Raman spectra requires sophisticated computing techniques due to their tremendous complexity and thousands of variables. Because they preserve correlations between observed process variables and spectral information while reducing spectral complexity, chemometric techniques—particularly partial least squares (PLS) regression—have long been utilized for quantitative analysis [5]–[8].

Despite the success of chemometric models, complex biological processes can entail nonlinear interactions that linear techniques might not adequately reflect. The extraction of hidden associations from high-dimensional spectral datasets has been made possible by recent developments in machine learning. In analytical and biological applications, machine learning algorithms, including Support Vector

Regression, Random Forest, gradient boosting techniques, and neural networks, have shown promise for increasing prediction accuracy [9]–[12].

Among advanced machine learning approaches, XGBoost has gained significant attention because of its ability to efficiently model nonlinear relationships and handle complex datasets [13].

Recently, machine learning has also been successfully applied to spectroscopic Process Analytical Technology (PAT) for pharmaceutical manufacturing. For example, Fiorenza et al. demonstrated that combining near-infrared (NIR) spectroscopy with machine learning models enabled accurate prediction of residual moisture during lyophilization, highlighting the potential of artificial intelligence to support real-time quality assurance in regulated pharmaceutical processes [14].

However, problems with model transparency are frequently brought about by improved predictive capability. Understanding why a model produces a particular forecast is crucial for regulatory approval in regulated production settings. By determining how each variable contributes to predictions, explainable artificial intelligence (XAI) techniques like SHAP (SHapley Additive exPlanations) offer ways to interpret machine learning conclusions [15], [16]. Artificial intelligence-based industrial decisions may be more confident if explainability is incorporated into PAT systems.

The development of reliable and comprehensible predictive models for Raman-based bioprocess monitoring still faces significant obstacles, despite notable advancements in spectroscopy-assisted machine learning for Process Analytical Technology (PAT). Previous studies have demonstrated the value of Raman spectroscopy, chemometric modeling, and machine learning-assisted PAT for improving process monitoring and predictive performance [17]–[19]. Further demonstrating the expanding role of artificial intelligence in spectroscopy-based PAT systems, Fiorenza et al. recently showed how machine learning-integrated near-infrared (NIR) spectroscopy can be successfully applied for residual moisture prediction during pharmaceutical lyophilization [14]. For Raman spectroscopy-based prediction of multiple fermentation metabolites, relatively few studies have systematically compared conventional chemometric methods with sophisticated machine learning algorithms, incorporating explainable artificial intelligence to enhance model transparency and regulatory confidence.

Thus, utilizing Raman spectroscopy data from Escherichia coli fermentation, this study creates a comparable predictive modeling framework. For the prediction of glucose and acetate, several machine learning techniques are assessed, and SHAP-based explainable artificial intelligence is integrated to find significant spectrum characteristics influencing model predictions. This work advances the wider application of PAT and Real-Time Release Testing (RTRT) in next-generation bioprocess manufacturing by expanding machine learning-assisted spectroscopy beyond pharmaceutical lyophilization to Raman-based microbial fermentation monitoring.

### 1.1. Research Question

Primary Research Question: Which machine learning model provides the most accurate and dependable prediction of critical bioprocess parameters from Raman spectroscopic data to enable Process Analytical Technology (PAT) and Real-Time Release Testing (RTRT)?

Goal of the Research: The goal of this research is to create and evaluate an explainable machine learning system that can use Raman spectral data to predict significant fermentation parameters. In order to ascertain if model selection should be based on the biological properties of certain process variables, the study contrasts sophisticated artificial intelligence models with conventional chemometric techniques.

Specifically, this research evaluates:

1. Whether Raman spectroscopy combined with machine learning can accurately predict glucose and acetate concentrations during fermentation.
2. Whether advanced artificial intelligence models outperform traditional chemometric methods.
3. Which Raman spectral features contribute most significantly to model predictions using explainable AI methods?

## 2. Literature Review

### 2.1. Raman Spectroscopy in Bioprocess Monitoring

Raman spectroscopy has become an important analytical technology for biomanufacturing because it enables continuous monitoring without disrupting the production process. Several studies have demonstrated that Raman-based models can accurately estimate important fermentation variables, including nutrient concentrations and metabolic byproducts [20], [21].

Raman spectroscopy has advantages over traditional analytical techniques, including quick measurement, little sample preparation, and compatibility with automated manufacturing processes [22]. Because of these features, Raman spectroscopy is an excellent choice for implementing Process Analytical Technology (PAT) in contemporary biopharmaceutical manufacturing settings [18].

### 2.2. Chemometric Approaches for PAT Applications

Spectroscopic data analysis has historically been based on chemometric techniques [7], [8], [23]. Because it can handle highly correlated spectrum variables and provide quantitative correlations between Raman signals and biological data, partial least squares (PLS) regression is still one of the most popular techniques [17] [18].

PLS-based approaches have been successfully applied in pharmaceutical and biological spectroscopy due to their ability to reduce spectral dimensionality while maintaining important information related to process variables [23].

However, because biological systems are intrinsically complex, nonlinear metabolic interactions may not always be captured by linear models. This constraint has prompted researchers to investigate machine learning techniques that can predict more complex patterns [6].

### 2.3. Machine Learning for Raman-Based Bioprocess Prediction

Machine learning algorithms provide new opportunities for analyzing complex Raman datasets. Support Vector Regression has demonstrated strong performance in high-dimensional analytical problems because of its ability to model nonlinear relationships and handle complex feature spaces [10].

By merging several decision models, ensemble learning techniques like Random Forest and XGBoost enhance prediction performance. While XGBoost has shown outstanding performance in numerous scientific applications due to its computational efficiency and predictive power, Random Forest has been frequently employed for robust classification and regression tasks [11], [13].

The promise of artificial intelligence and machine learning techniques for bioprocess optimization, quality prediction, and real-time manufacturing control has also been emphasized by recent studies [24].

Recent research has investigated the integration of machine learning with various spectroscopic modalities for Process Analytical Technology, in addition to Raman spectroscopy. Fiorenza et al. demonstrated the increasing significance of artificial intelligence for real-time process monitoring and release testing by reporting that NIR spectroscopy in conjunction with ensemble machine learning models achieved accurate monitoring of residual moisture during pharmaceutical lyophilization [14]. These results highlight the necessity to look into modality-specific performance in bioprocessing applications while supporting the wider use of spectroscopy-driven machine learning in pharmaceutical manufacture.

### 2.4. Explainable Artificial Intelligence for PAT

Despite their high accuracy, machine learning models need to be transparent and easy to understand in order to be used in pharmaceutical manufacturing. By identifying important factors and interactions, explainable AI techniques offer insight into model decision-making [25].

By enhancing the comprehension of spectral variables influencing model predictions, the integration of explainable AI with Raman spectroscopy may boost regulatory confidence and hasten the implementation of AI-driven PAT systems [26].

**Table 1.** Summary of Machine Learning Approaches Used for Raman-Based Bioprocess Prediction. Comparison of machine learning algorithms evaluated for Raman spectroscopy-based bioprocess monitoring.

| Model | Category | Main Advantage | Application in Spectroscopy |
| --- | --- | --- | --- |
| PLS Regression | Chemometric model | High interpretability and regulatory acceptance [4] | Quantitative Raman calibration |
| SVR | Machine learning | Handles nonlinear relationships [4] | Spectral prediction problems |
| Random Forest | Ensemble learning | Feature importance analysis [11] | Complex biological datasets |
| XGBoost | Gradient boosting | High predictive performance [5] | Nonlinear process modeling |
| Neural Network | Deep learning | Learns complex patterns [8] | Advanced spectral analysis |

**Table 2.** Dataset Characteristics. Characteristics of the Raman spectroscopy dataset used for machine learning-based bioprocess parameter prediction.

| Parameter | Description |
| --- | --- |
| Biological system | *Escherichia coli* fermentation |
| Analytical method | Raman spectroscopy |
| Number of Raman spectra | 379 |
| Raman variables per spectrum | 1,870 |
| Prediction targets | Glucose and Acetate |
| Machine learning task | Regression |
| Input matrix dimension | $379 \times 1870$ |
| Target matrix dimension | $379 \times 2$ |

## 3. Methodology

### 3.1. Dataset Description

This study developed and assessed machine learning models for real-time bioprocess monitoring using the publicly accessible RamanBench Escherichia coli fermentation dataset. For the purpose of comparing analytical and machine learning techniques in biological and chemical applications, RamanBench offers standardized Raman spectroscopy datasets [19].

The selected dataset represents a microbial fermentation process in which Raman spectra were collected during *E. coli* cultivation. Glucose and acetate concentrations were selected as target variables because they represent important indicators of microbial metabolism and fermentation performance. Raman spectroscopy has previously demonstrated the capability to monitor important fermentation metabolites and process variables in biological systems [20], [21]. The dataset characteristics are summarized in Table 2.

### 3.2. Raman Spectral Preprocessing

Raw Raman spectra frequently contain unwanted variation caused by fluorescence background, instrumental noise, and differences in sample measurement conditions. Appropriate preprocessing is therefore required before predictive modeling because spectral quality strongly influences chemometric and machine learning performance [17], [22].

The preprocessing workflow consisted of three major steps.

- **Spectral Smoothing:** Savitzky–Golay filtering was applied to reduce random noise while maintaining Raman peak characteristics. This approach is widely used in spectroscopy because it improves signal quality without significantly changing spectral information [18].
- **Standard Normal Variate Transformation:** Standard Normal Variate (SNV) normalization was applied to reduce scattering effects and intensity variation between spectra. SNV is commonly used in chemometric analysis of spectroscopic datasets to improve model robustness and reproducibility [5].
- **Feature Scaling:** Spectral variables were standardized before model training to ensure that machine learning algorithms were not influenced by differences in variable magnitude.

After preprocessing, the dataset contained no missing values and was used for model development.

### 3.3. Machine Learning Framework

Five regression algorithms were evaluated to compare traditional chemometric approaches with advanced artificial intelligence methods.

Five regression algorithms were evaluated to compare traditional chemometric approaches with advanced artificial intelligence methods.

- Partial Least Squares Regression (PLS): Because PLS regression is still often utilized in Raman-based PAT applications, it was chosen as the conventional chemometric benchmark. By creating latent variables that maximize covariance between spectral predictors and process parameters, the technique lowers spectral dimensionality [5], [8], [10].
- Support Vector Regression (SVR): SVR was used because it can apply kernel-based adjustments to capture nonlinear correlations between Raman spectral characteristics and biological data. In high-dimensional analytical datasets, SVR has proven to perform well [9], [10].
- Random Forest Regression: Because Random Forest mixes several decision trees to increase prediction stability and decrease overfitting, it was chosen as an ensemble learning technique. Furthermore, Random Forest offers changeable significance information that is helpful for spectral interpretation [11].
- Extreme Gradient Boosting (XGBoost): XGBoost was included as an advanced gradient boosting algorithm. The method improves prediction performance by sequentially correcting errors from previous decision trees and has demonstrated excellent performance in complex data modeling applications [13].
- Neural Network Regression: A multilayer perceptron neural network was evaluated to investigate whether deep learning approaches could extract complex spectral patterns. Neural networks have been increasingly applied to spectroscopy because of their ability to learn nonlinear relationships from high-dimensional datasets [12].

### 3.4. Model Evaluation

Model performance was evaluated using three commonly reported regression metrics:

- Coefficient of determination (R²): Measures the proportion of variance explained by the model.
- Root Mean Square Error (RMSE): Evaluates average prediction error magnitude.
- Mean Absolute Error (MAE): Measures average absolute difference between predicted and experimental values.

Higher R² and lower RMSE/MAE indicate better predictive performance. These evaluation metrics are commonly used for assessing quantitative prediction models in spectroscopy-based analytical applications [17], [18].

### 3.5. Explainable Artificial Intelligence Analysis

To improve model interpretation, SHAP analysis was applied to identify Raman spectral variables contributing to model predictions. SHAP values provide an estimate of how individual features influence machine learning outputs and have been increasingly used for interpreting complex AI models [15], [16].

The integration of explainable AI allows the developed framework to move beyond prediction accuracy by providing information about important Raman spectral regions associated with biological changes. This approach supports greater transparency and may improve acceptance of AI-driven PAT systems in regulated biomanufacturing environments [25], [26].

## 4. Results and Discussion

### 4.1. Comparative Model Performance for Glucose Prediction

The performance of all models was evaluated for glucose and acetate prediction. The predictive capability of Raman-based machine learning models was assessed using the coefficient of determination (R²), root mean square error (RMSE), and mean absolute error (MAE). These metrics are commonly used for evaluating quantitative spectroscopy models because they provide information regarding both explained variance and prediction error magnitude [17], [18].

Similar observations have been reported by Fiorenza et al., where chemometric and machine learning models demonstrated strong predictive capability for spectroscopic quality attribute prediction in pharmaceutical manufacturing. Their findings suggest that model performance depends on both the analytical technique and the physicochemical characteristics of the target variable [14].

**Table 3.** Machine Learning Model Performance for Glucose and Acetate Prediction. Performance comparison of machine learning models for Raman-based prediction of glucose and acetate concentrations.

| Target | Model | R<sup>2</sup> | RMSE | MAE |
| --- | --- | --- | --- | --- |
| Glucose | PLS Regression | 0.972 | 0.216 | 0.175 |
| Glucose | SVR | 0.954 | 0.279 | 0.205 |
| Glucose | Random Forest | 0.898 | 0.416 | 0.253 |
| Glucose | XGBoost | 0.918 | 0.374 | 0.215 |
| Glucose | Neural Network | 0.831 | 0.535 | 0.367 |
| Acetate | PLS Regression | 0.500 | 0.161 | 0.121 |
| Acetate | SVR | 0.339 | 0.186 | 0.131 |
| Acetate | Random Forest | 0.712 | 0.123 | 0.086 |
| Acetate | XGBoost | 0.722 | 0.120 | 0.081 |
| Acetate | Neural Network | -1.290 | 0.345 | 0.219 |

**Figure 1.**
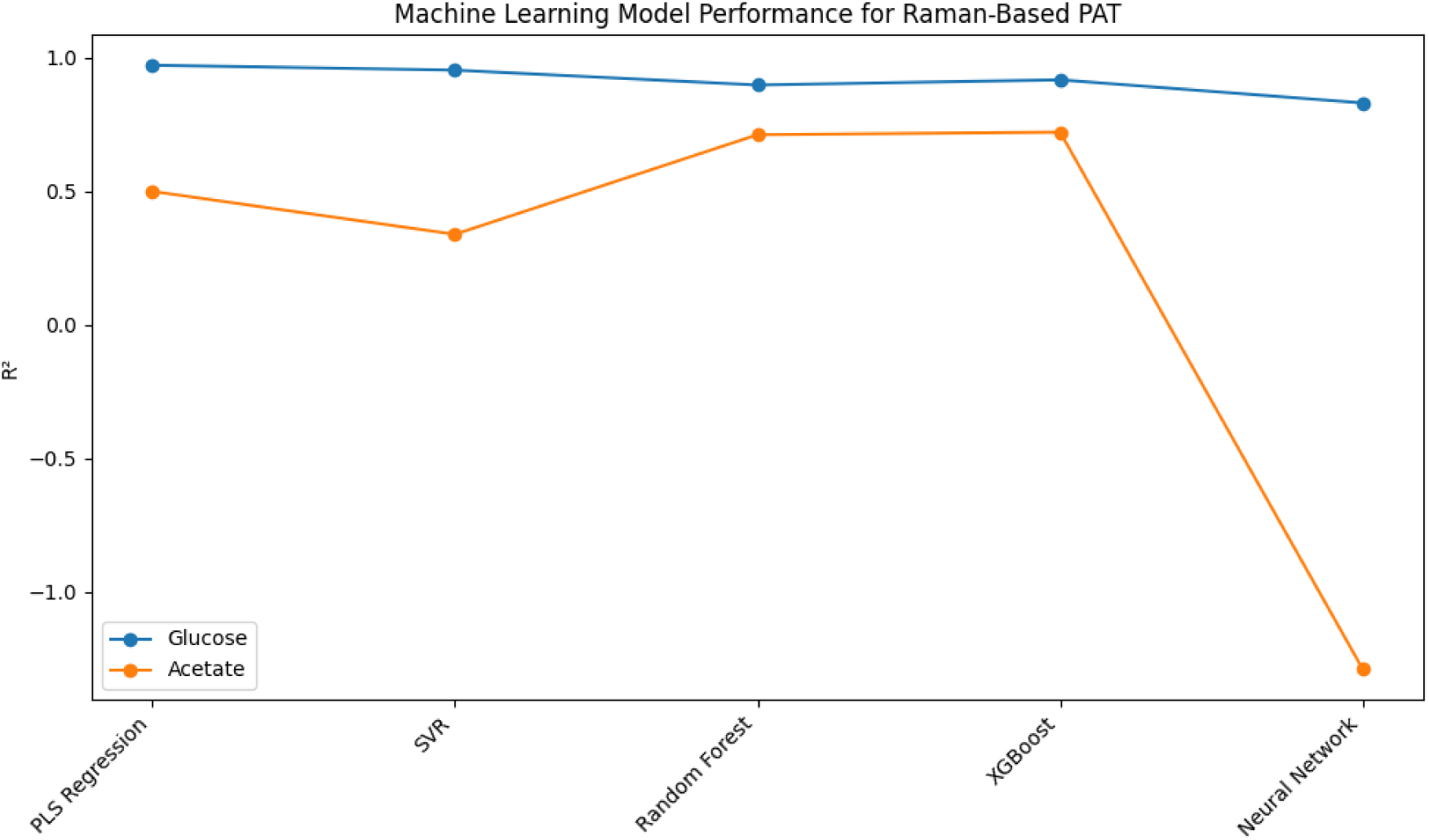
Comparative performance of machine learning models based on coefficient of determination (R²), RMSE, and MAE values. PLS regression demonstrated the highest accuracy for glucose prediction, whereas XGBoost achieved superior acetate prediction performance.

### 4.2. Glucose Prediction Performance

PLS regression achieved the highest glucose prediction accuracy with an R² value of 0.972. This result demonstrates that glucose concentration showed strong relationships with Raman spectral characteristics.

The strong performance of PLS is consistent with previous studies showing that chemometric models remain highly effective for predicting well-characterized analytes in bioprocess environments [5], [8], [10]. PLS-based models have historically been successful in Raman applications because they can reduce spectral complexity while maintaining information associated with important process variables.

Despite offering more flexibility, sophisticated models like XGBoost and neural networks did not increase the accuracy of glucose prediction. This suggests that higher model complexity does not always translate into better performance.

Because of their ease of use, interpretability, and regulatory acceptance, classic chemometric techniques may continue to be preferred for parameters with reasonably stable spectral relationships. This finding is consistent with other PAT research showing that transparent and well-characterized models are still useful for applications in pharmaceutical manufacturing [5], [18].

### 4.3. Comparative Model Performance for Acetate Prediction

Acetate prediction demonstrated a different modeling behavior.

| Model | $R^2$ | RMSE | MAE |
| --- | --- | --- | --- |
| XGBoost | 0.722 | 0.120 | 0.081 |
| Random Forest | 0.712 | 0.123 | 0.086 |
| PLS Regression | 0.500 | 0.161 | 0.121 |
| SVR | 0.339 | 0.186 | 0.131 |
| Neural Network | -1.290 | 0.345 | 0.219 |

**Figure 2.**
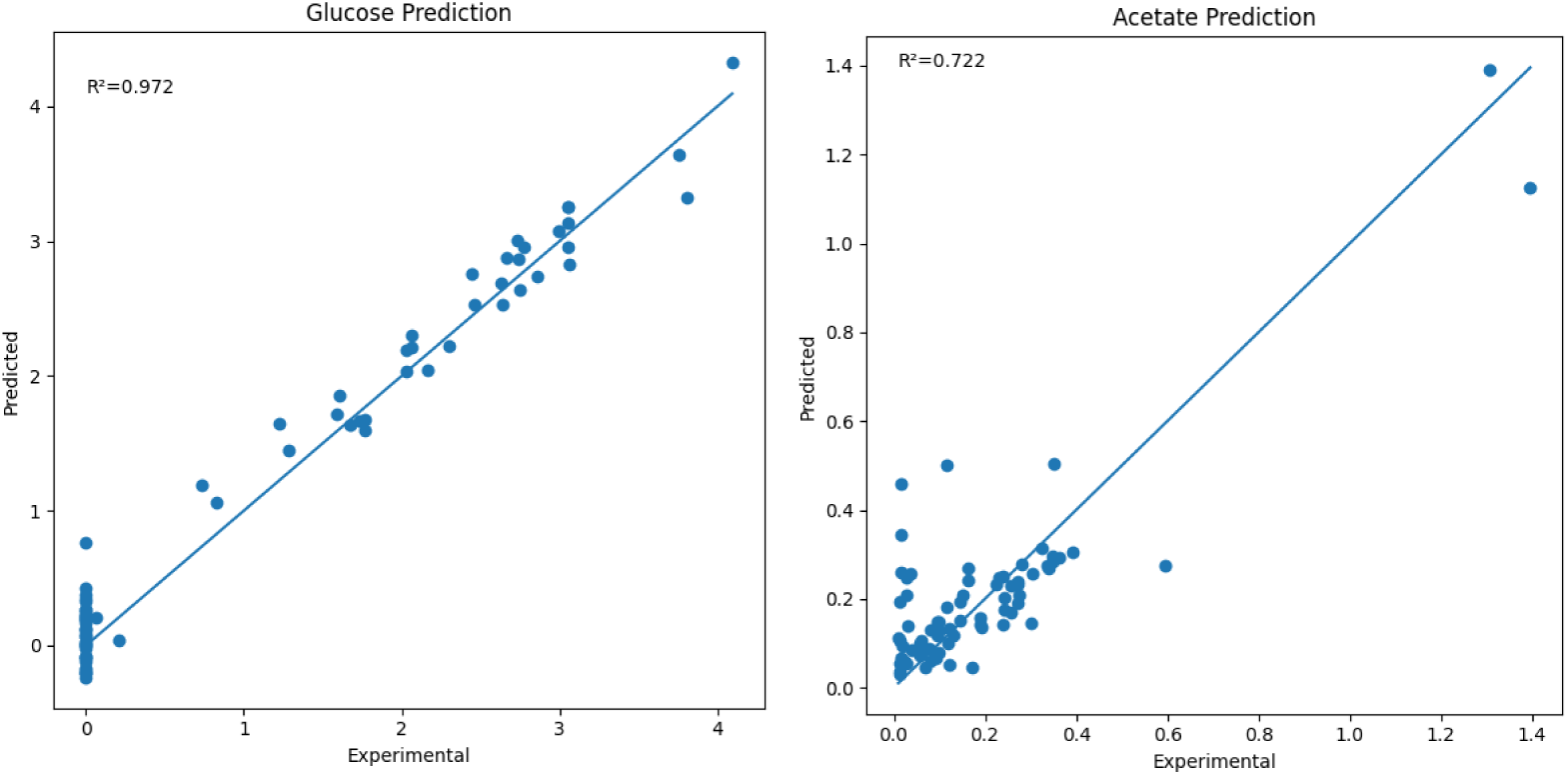
Experimental versus predicted concentrations for Raman-based machine learning models. (A) Glucose prediction using PLS regression. (B) Acetate prediction using XGBoost regression.

Unlike glucose prediction, acetate concentration was better predicted using nonlinear ensemble models.

XGBoost achieved the highest predictive accuracy, suggesting that acetate formation involves more complex spectral relationships that cannot be completely represented through linear latent variables. Gradient boosting approaches such as XGBoost are capable of modeling nonlinear interactions and complex feature relationships, making them suitable for biological datasets [13].

Acetate accumulation during microbial fermentation depends on metabolic regulation, carbon overflow pathways, and environmental conditions. These biological interactions may generate nonlinear relationships between Raman spectral signals and acetate concentration [22], [24].

The difference between glucose and acetate prediction demonstrates that bioprocess monitoring systems should not rely on a single universal modeling strategy. Instead, model selection should consider the biochemical characteristics of each process parameter.

The improved performance of XGBoost demonstrates that nonlinear machine learning methods can provide advantages for complex biological parameters [11], [13].

Unlike the pharmaceutical freeze-drying application investigated by Fiorenza et al. [14], the present study demonstrates that model selection is also influenced by metabolite-specific nonlinear behavior in microbial fermentation, with XGBoost outperforming conventional chemometric models for acetate prediction.

### 4.4. Biological and PAT Implications

The results provide important implications for future RTRT implementation.

A common assumption is that artificial intelligence models will always outperform traditional chemometric methods. However, this study demonstrates a more balanced conclusion.

**Figure 3.**
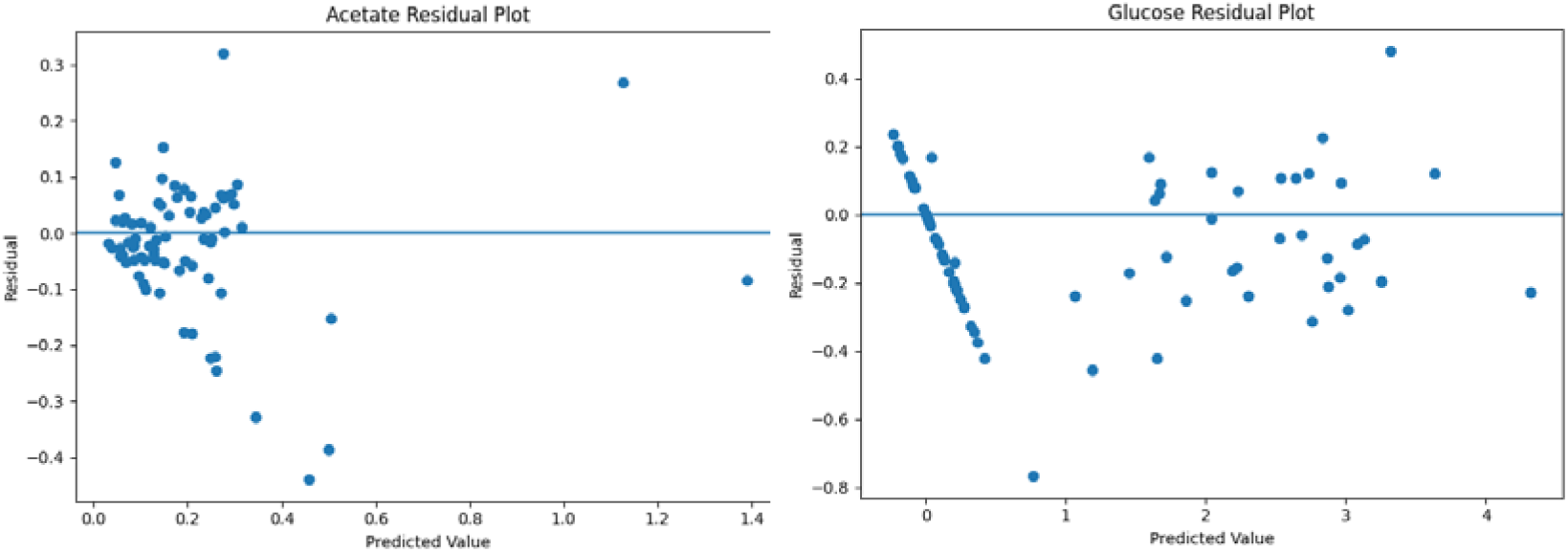
Residual analysis of Raman-based prediction models. Residual distributions demonstrate prediction reliability and absence of systematic errors across the concentration range.

For relatively predictable analytes such as glucose:

- PLS provides excellent performance.
- The model is computationally simple.
- Regulatory interpretation is easier.

For complex metabolic products such as acetate:

- Nonlinear machine learning approaches provide improved prediction.
- Advanced models capture hidden spectral relationships.

Therefore, a hybrid PAT strategy may provide the most effective approach: Chemometric models for well-characterized parameters + AI models for complex biological behaviors.

This approach is in line with current developments in biopharmaceutical manufacturing, where integrating artificial intelligence with well-established analytical techniques is thought to be a viable way to enhance process knowledge and control [24], [26].

The results of this study and earlier research by Fiorenza et al. [14] show that machine learning-enhanced spectroscopy has a wide range of applications in the pharmaceutical and biopharmaceutical manufacturing industries. Raman spectroscopy showed complementary strengths for real-time monitoring of fermentation metabolites, underscoring the adaptability of spectroscopic-driven PAT solutions, while NIR spectroscopy was found to be beneficial for monitoring residual moisture during lyophilization.

### 4.5. Explainable AI Contribution

The integration of SHAP analysis provides an additional layer of interpretation beyond predictive accuracy.

Rather than treating machine learning models as black boxes, SHAP identifies Raman spectral regions contributing to predictions. This improves confidence in AI-based PAT systems by linking model decisions with measurable biochemical information.

SHAP-based interpretation methods provide feature-level explanations by estimating the contribution of individual variables to model predictions and have become increasingly important for improving the transparency of machine learning systems [15], [16].

Such interpretability is particularly important for pharmaceutical manufacturing because regulatory acceptance requires understanding of model behavior, robustness, and reliability [25], [26].

**Figure 4.**
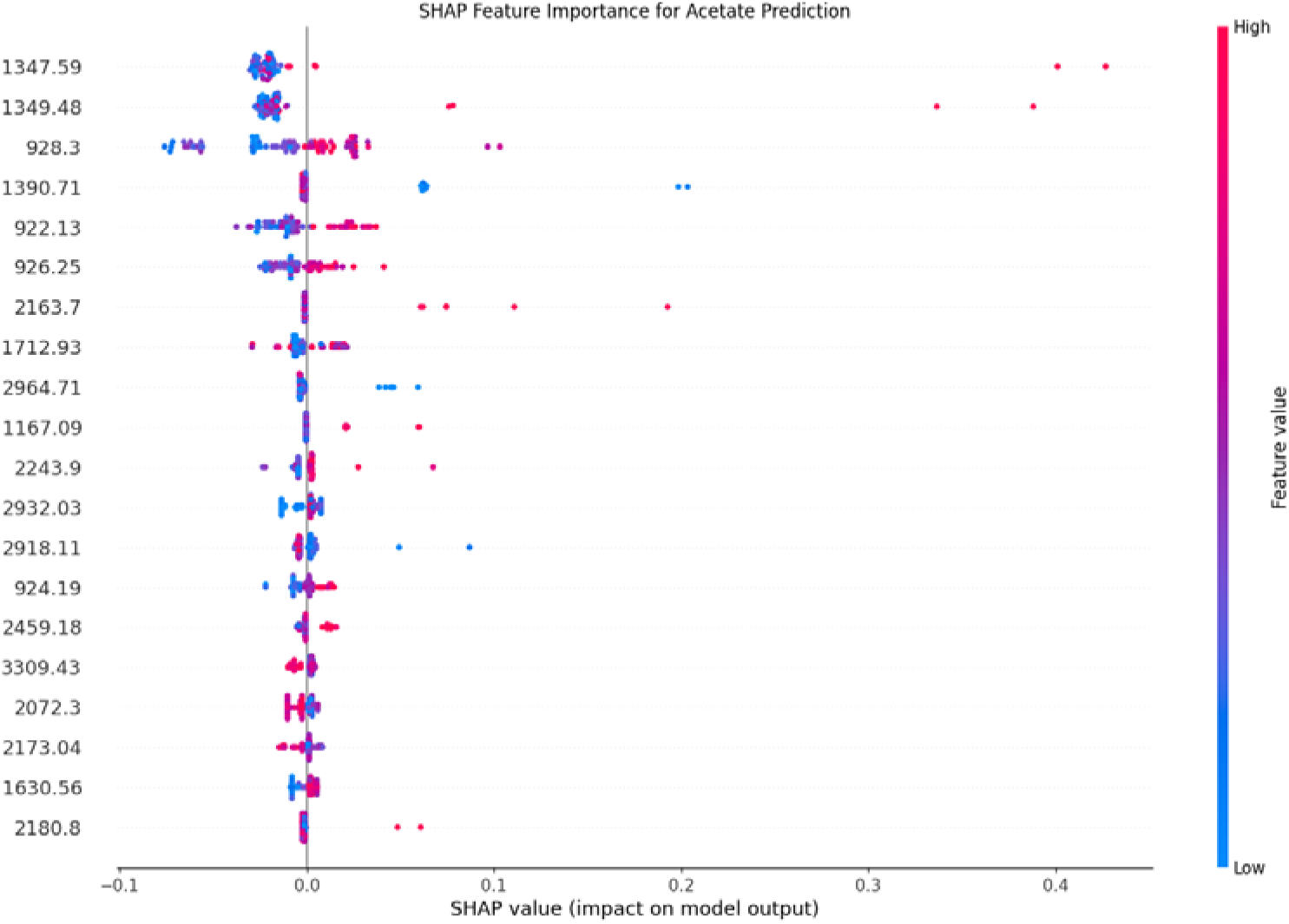
SHAP-based interpretation of Raman spectral features contributing to model prediction. Important Raman regions were identified to improve model transparency and provide potential biochemical interpretation.

## 5. Advantages of the Proposed Framework

The proposed framework provides several advantages for next-generation bioprocess manufacturing.

- In addition to facilitating real-time process monitoring, Raman spectroscopy reduces reliance on offline sampling and laboratory analysis by enabling continuous, quick, and non-destructive assessment of bioprocess variables [1], [2], [3], [20], [23].
- Combining multiple machine learning approaches allows selection of the most appropriate predictive model according to the biochemical characteristics of each process parameter. The comparative evaluation demonstrated that traditional chemometric models remain highly effective for analytes with relatively linear spectral relationships, whereas nonlinear ensemble learning methods provide improved performance for more complex biological responses [10], [11], [13], [24].
- The integration of explainable artificial intelligence enhances confidence in machine learning predictions by identifying the Raman spectral regions that contribute most strongly to model outputs rather than treating prediction models as black boxes [15], [16], [25].
- By facilitating automated process monitoring, quick decision-making, and enhanced factory control, the suggested architecture enables the future application of Process Analytical Technology (PAT) and Real-Time Release Testing (RTRT). Next-generation digital biomanufacturing systems must have these features [6], [10], [23], [26].
- By extending machine learning-assisted process monitoring to Raman spectroscopy and microbial fermentation, the suggested framework enhances earlier spectroscopy-based PAT systems documented for pharmaceutical manufacture [14].

## 6. Limitations

Despite the promising results obtained in this study, several limitations should be considered.

- A single *Escherichia coli* fermentation dataset was used in the study. Before industrial deployment, further validation utilizing numerous microbial strains, mammalian cell cultures, and large-scale production platforms is required, even though the RamanBench dataset offers realistic Raman spectra for assessing predictive models [19], [20], [23].
- Only 379 Raman spectra were included in the dataset, which is quite few when compared to datasets usually needed for deep learning applications. This restriction probably played a part in the neural network model’s comparatively subpar performance, especially when it came to acetate prediction. Deep learning performance and generalizability would probably be enhanced by larger datasets gathered from various fermentation processes [12], [24].
- Predictive accuracy was the main objective of this study, not complete industrial deployment. Future research should assess the resilience of the model under realistic manufacturing settings, such as batch-to-batch variation, sensor drift, process disruptions, and instrument variability, all of which can affect forecast reliability [5], [10], and [11].
- Before routine industrial adoption can be accomplished, regulatory implementation of AI-based PAT systems necessitates additional validation processes, such as calibration transfer, model lifecycle management, continuous performance monitoring, and compliance with pharmaceutical quality guidelines [1], [26].
- In order to enable complete Process Analytical Technology spanning upstream and downstream biopharmaceutical manufacturing, future research should examine the integration of various spectroscopic techniques, including Raman and NIR spectroscopy, within unified machine learning frameworks [14].

## 7. Future Work

Future research should focus on extending the proposed framework toward industrial-scale implementation and intelligent biomanufacturing. Potential directions include:

1. To increase the robustness and generalizability of the model, continuous bioreactor datasets gathered from various manufacturing facilities and operating conditions are integrated [11], [26].
2. Development of hybrid physics-informed machine learning models that combine mechanistic fermentation knowledge with data-driven prediction techniques to improve both prediction accuracy and model interpretability [24], [26].
3. Real-time deployment of predictive models using edge computing systems directly connected to Raman spectrometers for continuous online monitoring and automated process control [10], [23].
4. Integration of additional Process Analytical Technology sensors, including pH, dissolved oxygen, biomass, conductivity, and metabolite analyzers, to develop multi-sensor predictive frameworks for comprehensive process monitoring [2], [23].
5. To promote industrial acceptance, regulatory-ready AI processes with automated model monitoring, drift detection, recalibration techniques, and quality assurance protocols are being developed [25], [26].
6. Expansion of explainable artificial intelligence methods to identify biochemical mechanisms associated with Raman spectral changes and improve biological interpretation of machine learning predictions [15], [16], [25].

## 8. Conclusion

A machine learning-enabled Raman spectroscopy framework for Process Analytical Technology (PAT) and Real-Time Release Testing (RTRT) applications in bioprocess manufacturing was created and assessed in this study.

The findings showed that the biochemical properties of the target analyte have a significant impact on prediction performance. Excellent glucose prediction performance was attained using partial least squares regression, suggesting that traditional chemometric techniques are still quite successful for analytes with comparatively linear spectral correlations. On the other hand, XGBoost offered better acetate concentration prediction, proving that nonlinear ensemble learning algorithms may more successfully mimic intricate biological processes [10], [13].

A possible route toward transparent, comprehensible, and automated bioprocess monitoring is the combination of explainable artificial intelligence, machine learning, and Raman spectroscopy. Advanced machine learning techniques should be viewed as complementing technologies that can handle nonlinear biochemical interactions that are challenging to capture using classic statistical methods, rather than as a replacement for conventional chemometric techniques [15], [16], [24], [25].

Together with recent advances in NIR spectroscopy-based PAT for pharmaceutical lyophilization [14], the present work demonstrates that spectroscopy integrated with machine learning has the potential to support intelligent, real-time monitoring across diverse biopharmaceutical manufacturing processes. The complementary findings of both studies reinforce the expanding role of artificial intelligence in enabling robust Process Analytical Technology and Real-Time Release Testing.

Overall, by integrating explainable artificial intelligence, predictive analytics, and spectroscopic sensing into a single decision-support system, the suggested framework advances intelligent biomanufacturing. In biopharmaceutical production, the framework has the potential to enhance process comprehension, decrease analytical delays, bolster regulatory confidence, and ease the deployment of next-generation automated manufacturing platforms that can provide dependable real-time quality decisions [14], [24]–[26].

## Acknowledgment

Grammarly was used for clarity, but the authors take full responsibility for the content, including all research, analysis, and conclusions. The authors also used AI for Libraries, Functions, Graphs, and Tables.

